# Direct Pacbio sequencing methods and applications for different types of DNA sequences

**DOI:** 10.1101/2023.12.12.571020

**Authors:** Yusha Wang, Xiaoshu Ma, Lei Yang, Hua Ye, Ruikai Jia

## Abstract

The development of Sanger sequencing and next-generation sequencing methods within the past few years have assisted investigators profile the diversity and relative abundances of heterogenous species in vector preparations. Especially Recombinant adeno-associated viruses (rAAVs), genome editing, and mRNA related research are currently the most prominently investigated platform in different area and essentially use for synthetic biology, gene and cell therapy, food industrial and medicinal pharmer etc. area. However, these types of research related constructs always contain high GC sequences, poly structure, long-length DNA sequences and ITR repeats sequences.

Unfortunately, Sanger sequencing and NGS platforms may be inaccessible to investigators with limited resources, require large amounts of input material, or may require long wait times for sequencing and analyses. Recent advances with PacBio sequencing have helped to bridge the gap for quick and relatively inexpensive long-read sequencing needs. Specifically, long-read sequencing methods, like single molecule real-time (SMRT) sequencing, have been used to uncover truncations, chimeric genomes, and inverted terminal repeat (ITR) mutations in vectors. Recombinant adeno-associated virus (raav) is the most prominent platform in the field of current research, and its sequence is characterized by high GC, multi-structure, long sequence, genome, and repeat sequence. Sanger sequencing has certain defects in the detection of recombinant adeno-associated viruses. Meanwhile, Sanger needs to design sequencing primers based on known sequences to determine whether the sequences are correct. When sequence information is incomplete, it can only randomly design primers, obtain a sequence by luck, and then conduct the next round of sequencing. However, PacBio’s limitations and sample biases are not well-defined for sequencing. And sometimes the accuracy for base calling was low, resulting in a high degree of miscalled bases and false indels. These false indels led to read-length compression; thus, assessing heterogeneity based on read length is not advisable with current PacBio technologies. In this study, we explored the capacity for PacBio sequencing to directly interrogate content to obtain full-length resolution of encapsulated genomes. We found that the PacBio platform can cover the entirety of different type sequences like poly structure, long-length DNA fragment, high GC sequences and repeat sequences, especially the rAAV sequences from ITR to ITR without the need for pre-fragmentation. At the same time, the sequencing process was optimized to complete the sequencing of long difficult plasmids with the fewest plasmids and the fastest time. In summary, the optimization PacBio sequencing and novel bioinformation (BI) analysis method are able to correctly identify truncation hotspots in single-strand and self-complementary vectors using by SMRT sequencing and can serve as a rapid and low-cost alternative for proofing different type of sequences.

## Introduction

With the rapid development of sequencing technology, long-reads sequencing platforms, including Pacific Biosciences (PacBio) [1] and Oxford Nanopore Technologies (ONT) [2], are capable of end-to-end sequencing of entire cDNA molecules. Compared to next generation sequencing methods, these two platforms greatly increase the read length and can be used to solve various research problems.

The Single Molecule Real Time (SMRT) system developed by Pacific Biosciences applies the principle of synthesis while sequencing and uses the SMRT chip as the sequencing carrier. [3] Provides longer read lengths and faster running speeds. There are many ZMW holes in SMRT. ZMW (Zero-Mode Waveguides) hole [4]. Each SMRT Cell contains many these circular nanoholes, with a diameter of 50∼100nm. The holes use a zero-mode waveguide with a physical effect, and the outer diameter is smaller than the wavelength of the excited light. When DNA molecules enter the holes [5], the light emitted from the bottom of the excited light cannot penetrate the holes into the solution region above. Confined only to an area at the bottom that is large enough to cover the part of the DNA being detected, thereby collecting signals from that area, and minimizing background noise [6],

SMRTbell libraries are single-stranded hairpin junctions that connect long DNA molecules to sequencing junctions to form a stem-ring structure (and then add complementary sequencing primers and DNA polymerase molecules to the junctions) [7]. On-machine sequencing is to put the constructed library complex into the PacBio RS sequencer and load it into the nanopore of the sequencing chip SMRT Cell. DNA polymerase molecules are fixed at the bottom of the nanopore through covalent binding, usually one DNA molecule is fixed in each nanopore. Substrate dNTP and buffer solution required for DNA polymerization were added into the chip hole of SMRT Cell [8]. The four types of dNTP were equipped with four-color fluorescent labeling groups. According to the sequence of template chain nucleotides, the corresponding dNTP entered the DNA template chain, primer, and polymerase complex for chain extension reaction. At the same time, dNTP fluorescence signal was detected by zero-mode waveguide, fluorescence signal image was obtained, and DNA base sequence was obtained by calculation and analysis [9].

In terms of sequencing throughput, a SMRT Cell has 150,000 nanopore, and generally 1/3 ∼ 1/2 nanopore can generate effective DNA sequence reading, that is, 50,000 ∼ 70,000 effective DNA sequences can be obtained from each SMRT Cell [10]. PacBio’s CCS library performs multiple rounds of sequencing of a single fragment to improve accuracy. Since PacBio’s original errors are random errors, they can be corrected by themselves in CCS mode to improve data accuracy [11]. The average Read length of the most conservative Reads on the PacBio sequencing platform is 8-15kb. Assuming that the shortest Reads is 8Kb, it can also satisfy the requirement of correcting a 1.5Kb 16S full-length error for 5 times. According to official data, after the same fragment is sequenced for 4 times, the accuracy of a single read can reach at least 99% [12].

Gene synthesis is a method to synthesize double-stranded DNA in vitro, and it is one of the means to obtain the target gene. Sequencing is one of the most important methods to verify whether the synthesized genes are correct. Sanger sequencing is the most common plasmid sequencing method [13], and its principle is to add a certain proportion of labeled ddNTP to each of the four DNA synthesis reaction systems (including dNTP) (divided into: ddATP,ddCTP,ddGTP and ddTTP), through gel electrophoresis and autoradiography, can determine the DNA sequence of the molecule to be tested according to the position of the electrophoresis band [14]. Since the 2 ‘and 3’ of ddNTP do not contain hydroxyl groups, they cannot form phosphodiester bonds during DNA synthesis, so they can be used to interrupt DNA synthesis reactions. The following two conditions need to be met: 1) Template: that is, the sequence of recombinant plasmid is known, and all recombinant vectors are the same, and the sequence of inserted DNA fragments is unknown [15], which is exactly what we need to determine; 2) Primer: A nucleotide complementary to the plasmid sequence, and the complementary region should be selected at the junction with the unknown sequence DNA inserted into the plasmid. It can be synthesized in a DNA synthesizer using a chemical reaction [16], usually 17-25 bases. The primer is hybridized with the template by using temperature changes. If the synthesized gene contains complex structure, high/low GC, poly structure, Sanger sequencing effect may not be ideal [17]. In addition, each reaction of Sanger sequencing can only measure 600-800bp, and if the length of the synthetic gene is long, the sequencing cost will be greatly increased. In the sequencing process of PacBio, there was no PCR process, so Reads fragments with high GC content and low GC content were sequenced with similar probability [18], so GC Bias had little effect in third-generation sequencing. The library model of the PacBio data is a dumbbell structure with connectors at both ends, which is continuously sequenced around the library. The resulting sequence fragment is called polymerase reads [19], a sequence containing connectors, the length of which is determined by the activity of the reactive enzyme and the time spent on the machine [20]. At present, using the latest P6-C4 enzyme, the longest read length can reach more than 60kb. In this study, PacBio sequencing was applied to plasmid sequencing, and the application scope and accuracy of PacBio in plasmid sequencing were obtained through different types of plasmid sequencing.

## Result

### Large DNA-fragment sequencing by PacBio

Two 50kb and 32kb plasmids were constructed to test the effect of PacBio sequencing. The length of these two plasmids was measured by Agilent Bioanalyzer 2100 using g-tube (FIG. 1). The length of 32kb was 87%, ranging from 1,551bp to 13558bp, with an average length of 6894bp, accounting for 42.8% of all fragments. 83% of the length of 50kb breaks ranged from 1,927bp to 13514bp, with a mean length of 7815bp, accounting for 31.7% of all fragments (Figure 1B). The analysis of 32kb PacBio results showed (FIG. 2) that the distribution of 32kb plasmid showed that there were more reads in some places, but all of them were covered, indicating that all 32kb plasmid could be detected by PacBio (FIG. 2A). After BI analysis, it was concluded that the average sequencing depth of single base was 39, there were 6 loci with deletion, and the sequencing depth of 0 was deemed as deletion (Figure 2B). There were 12 base mutations, 8 of which had a mutation rate of 100%. There were three mutation sites with mutation rates of 98.28% (8949th base), 98.25% (8433rd base), and 96.55% (3077th base). Due to the error of PacBio machine, one other base was detected, and all mutation rates were not 100%. The mutation rate was 50% at site 17846, but the sequencing depth was only 2, so the mutation rate was not credible. Because the average sequencing depth of the plasmid was 39, but the sequencing depth of its site was 2, it could be determined that the deletion of its site (Figure 2C). There were four sites with insertion rates of 30%, which suggested that the clone was a mixture. To further determine if PacBio is suitable for sequencing long fragments, we constructed 50k plasmids for PacBio sequencing. According to the distribution of reads sub plasms, 3096 were deleted (Figure 3A) through BI analysis, and sanger sequencing showed that this plasmid was missing (Figure 3B). Seventeen of the bases were mutated, and three of the bases were not sequenced deep enough and should be considered missing (Table 1). With 32kb and 50kb plasmids, PacBio is friendly to long fragment sequencing.

**Figure 1.**
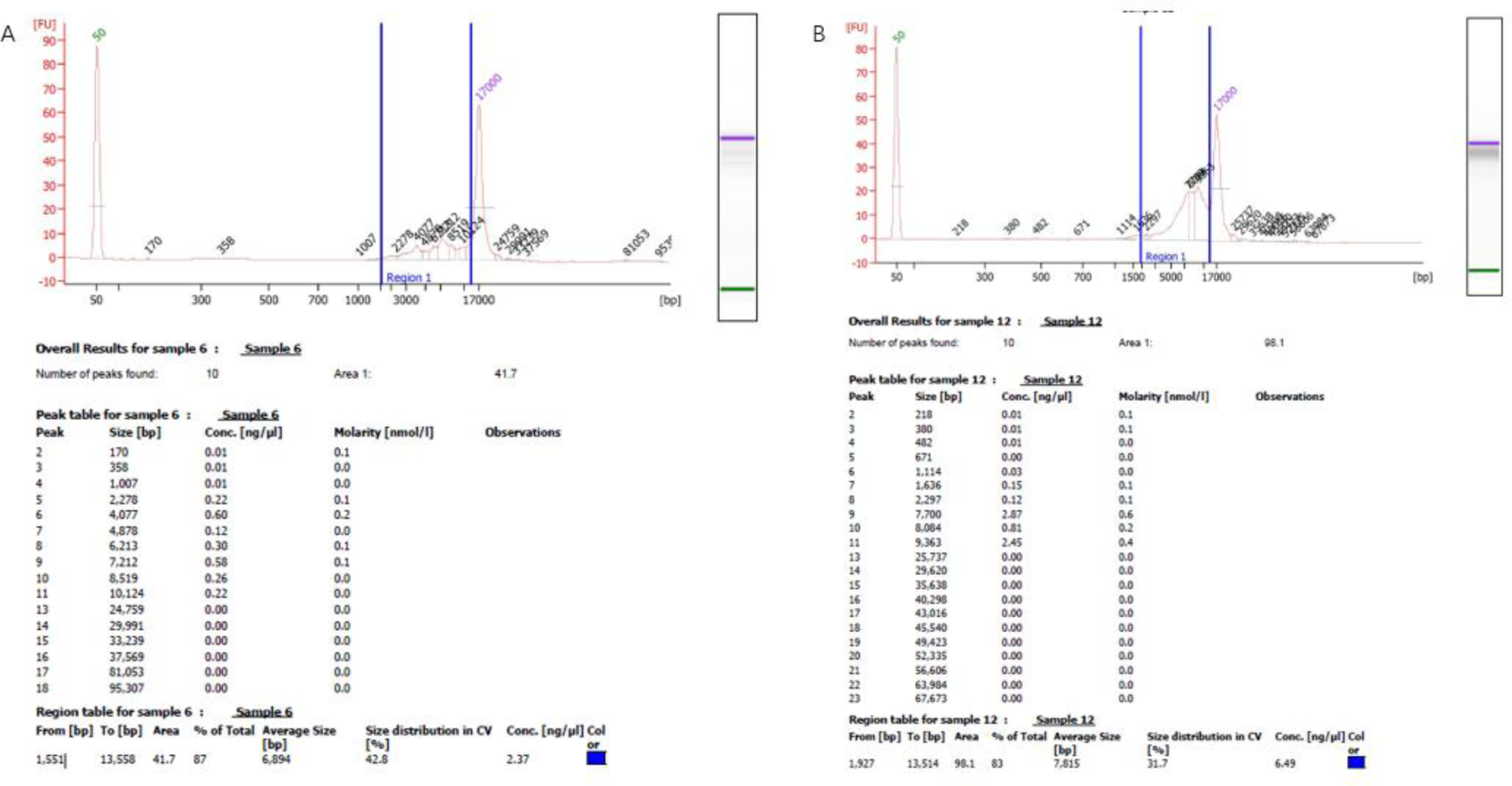
Agilent Bioanalyzer 2100 length determination.(A) The 23kb plasmid was interrupted by G-tube, and its average length was detected by Agilent Bioanalyzer 2100 after the interruption.(B) The 50kb plasmid was interrupted and its average length was measured. The length of the G-tube is not very uniform after the break, and the blue mark is the place where the fragment is most distributed, and the percentage of each length is marked. The Agilent Bioanalyzer 2100 machine calculates the average length based on the distribution of its fragment lengths.

**Figure 2.**
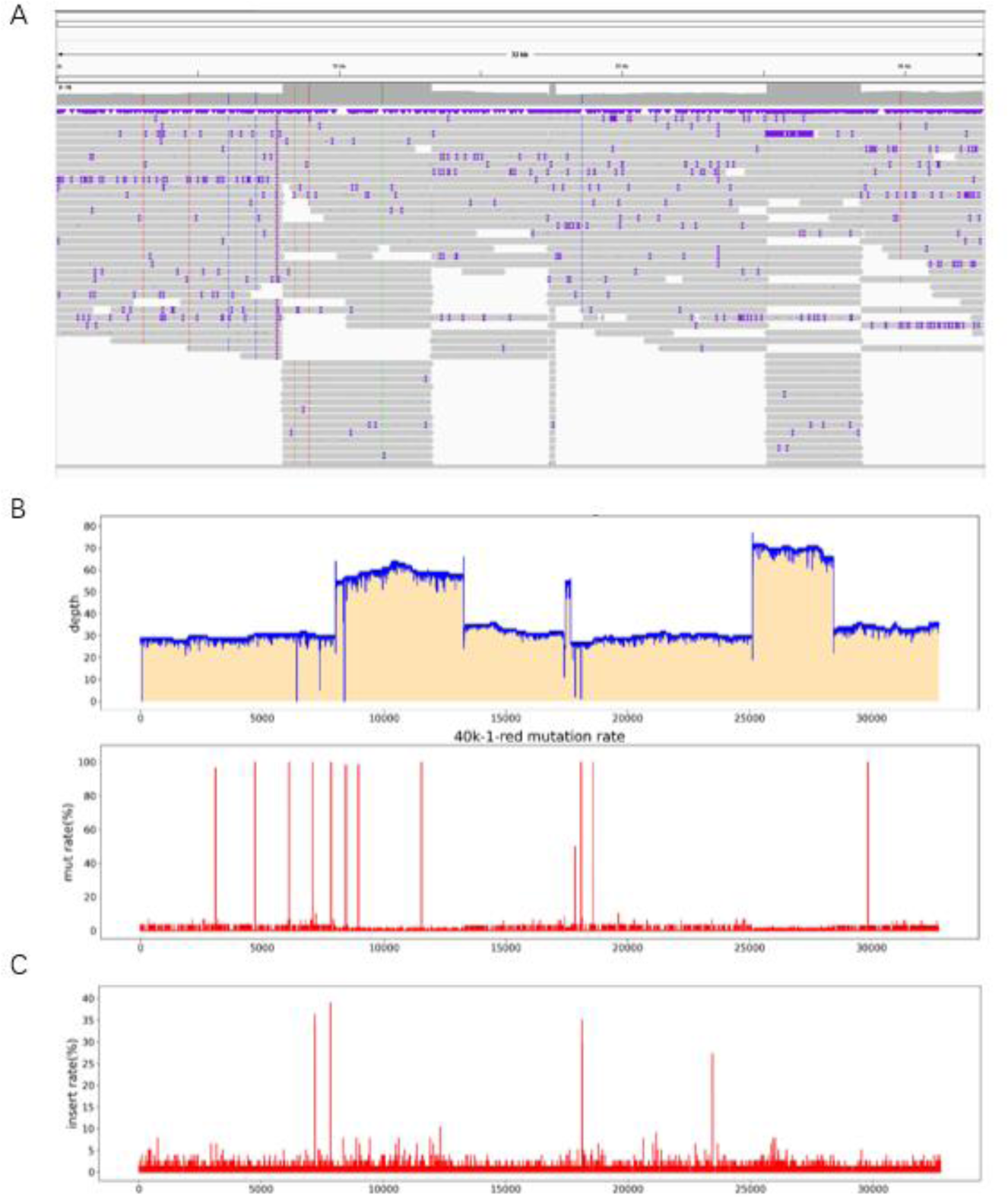
Sequencing results of 32kb plasmid PacBio. (A) The distribution of reads can be determined through IGV software. (B) Through BI analysis, the sequencing depth of each base is obtained. If the sequencing depth is 0, it is judged as missing. (C) Mutation rate at each site, with 11 sites having mutations.

**Figure 3.**
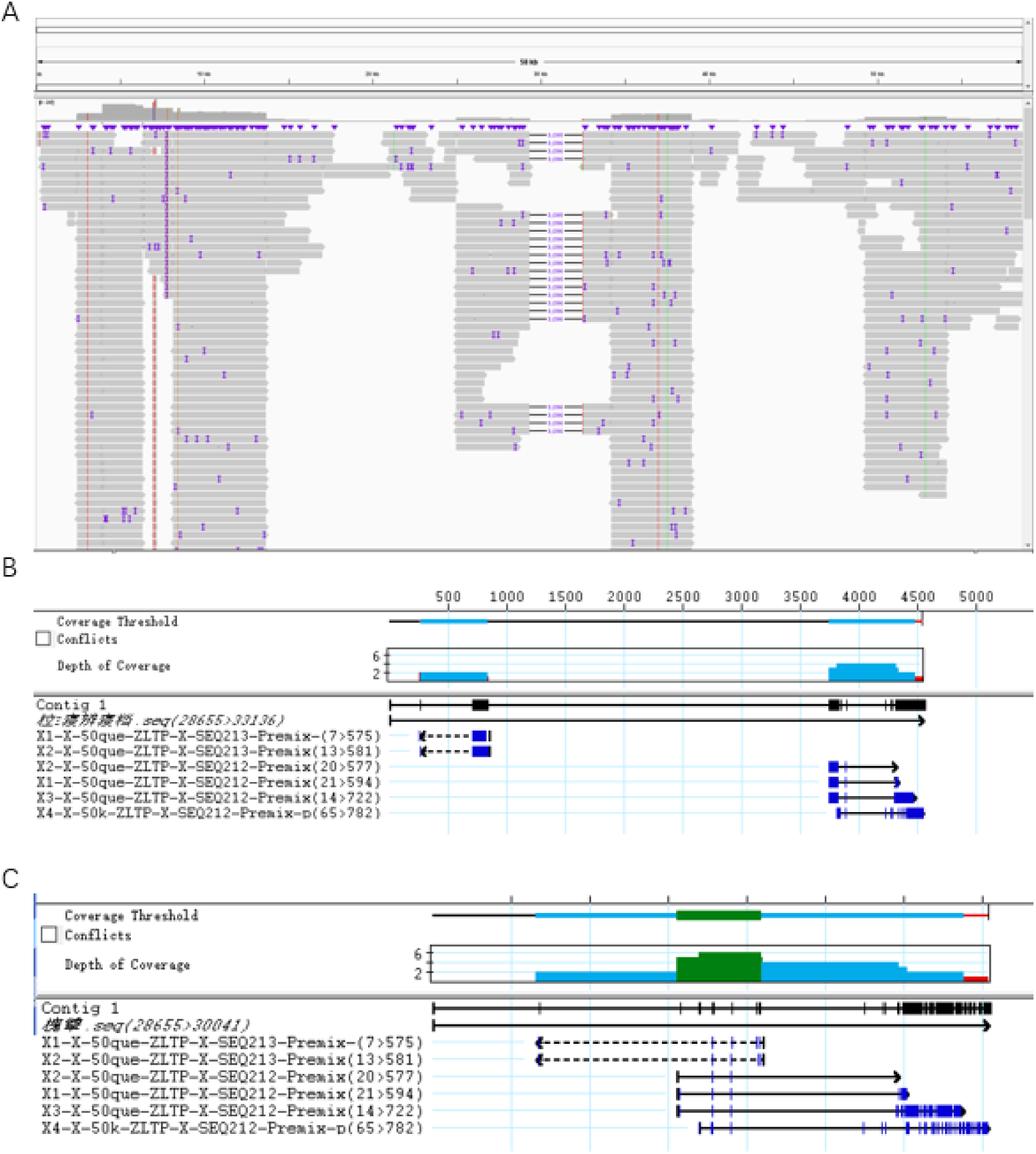
Distribution of plasmid reads of 50kb and sanger sequencing results. (A) The distribution of reads can be determined through IGV software. (B) Primers were designed at both ends of the deletion site with mutations at the tail end of both FR sanger results. (C) After removing the missing fragment, the remaining fragments were compared with the two Sangers, and the results were consistent.

**Table 1.** Sequencing results of PacBio mutant base of 50kb plasmid. . Pos represents the location of each base, Ref represents the original base, Depth represents the depth of sequencing, and ATCG represents the depth measured per base. Mut_rate(%) indicates the mutation rate.

| Id | Pos | Ref | Depth | A | C | G | T | Mut_rate(%) |
| --- | --- | --- | --- | --- | --- | --- | --- | --- |
| 50K_1_red | 6415 | A | 1 | 0 | 0 | 0 | 1 | 100 |
| 50K_1_red | 7828 | C | 149 | 0 | 0 | 149 | 0 | 100 |
| 50K_1_red | 7829 | T | 150 | 0 | 0 | 150 | 0 | 100 |
| 50K_1_red | 8433 | A | 161 | 0 | 0 | 161 | 0 | 100 |
| 50K_1_red | 15492 | A | 16 | 0 | 0 | 16 | 0 | 100 |
| 50K_1_red | 21273 | G | 4 | 4 | 0 | 0 | 0 | 100 |
| 50K_1_red | 32429 | A | 4 | 0 | 4 | 0 | 0 | 100 |
| 50K_1_red | 32431 | C | 4 | 0 | 0 | 4 | 0 | 100 |
| 50K_1_red | 32443 | G | 4 | 4 | 0 | 0 | 0 | 100 |
| 50K_1_red | 32444 | G | 5 | 4 | 1 | 0 | 0 | 100 |
| 50K_1_red | 32508 | C | 37 | 0 | 0 | 0 | 37 | 100 |
| 50K_1_red | 37519 | G | 58 | 58 | 0 | 0 | 0 | 100 |
| 50K_1_red | 42698 | G | 7 | 0 | 0 | 0 | 7 | 100 |
| 50K_1_red | 52858 | G | 19 | 19 | 0 | 0 | 0 | 100 |
| 50K_1_red | 3077 | C | 66 | 0 | 1 | 0 | 65 | 98.48 |
| 50K_1_red | 37005 | G | 56 | 0 | 0 | 1 | 55 | 98.21 |
| 50K_1_red | 25339 | G | 45 | 44 | 0 | 1 | 0 | 97.78 |
| 50K_1_red | 49287 | T | 2 | 0 | 1 | 0 | 1 | 50 |
| 50K_1_red | 902 | G | 35 | 0 | 0 | 23 | 12 | 34.29 |
| 50K_1_red | 57 | T | 4 | 1 | 0 | 0 | 3 | 25 |

### Poly structure sequencing by PacBio

In order to test the sequencing effect of PacBio on plasmids with poly structure, we constructed three plasmids with poly structure, namely polyA, polyC and polyG. The number of the three plasmids was 26 for A, 40 for C and 39 for G. After the interruption with g-tube, the library was built for PacBio sequencing. Through BI analysis, PacBio detected that when the number of polyA plasmid A was 26, the sequencing depth was 310 at most, while the sequencing depth of 25 A and 27 A was 160 and 155, thus it could be determined that the number of polyA of this plasmid was 26 (Figure 4A). The number of polyC plasmid C was 40, and the sequencing depth was 360 (Figure 4B). The number of polyG plasmid G was 39, and the sequencing depth was 950 (Figure 4C). Because of the instability of poly structure, in order to ensure the number of its preparation, the processing of the plasmid should be reduced during sequencing. In the process of building the database, PacBio uses physical methods to interrupt it, which can avoid the deletion or mutation of poly structure introduced by PCR reaction and ensure that the sequencing results match the reality to a certain extent. At the same time, PacBio belongs to single-molecule sequencing, which can measure most DNA molecules in a plasmid. According to the number of reads, it can effectively analyze whether the number of poly in the plasmid is single, and if not, which poly number exists respectively. However, sanger sequencing can only judge the quantity of most poly by the low peaks.

**Figure 4.**
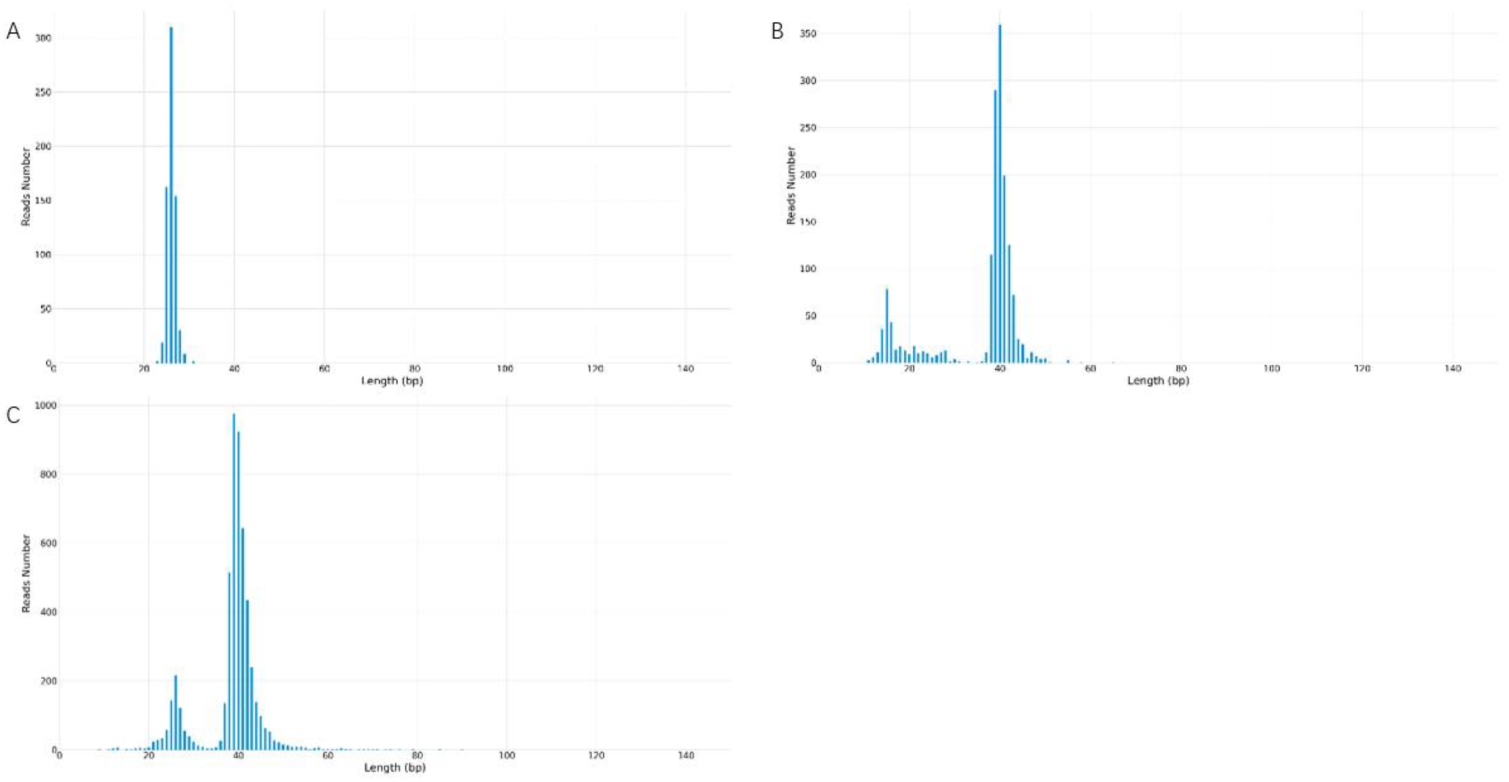
poly structure PacBio sequencing results. (A) PacBio sequencing results of polyA plasmid. The horizontal coordinate represents the length of A, and the vertical coordinate represents the sequencing depth of A. It can be seen that 26 A have the highest sequencing depth, and the number of polyA plasmid A detected by PacBio is 26. (B) PacBio sequencing results of polyC plasmid. The number of polyC plasmid C detected by PacBio was 40. (C) PacBio sequencing results of polyG plasmid. The number of polyG plasmid detected by PacBio was 40.

### Repeated structure sequencing by PacBio

In sanger sequencing, problems in sequencing results are caused by repetitive structures. The usual treatment is to design multiple sequencing primers and conduct positive and negative sequencing to ensure that the entire repetitive structure can be tested. In order to verify whether PacBio could detect the entire repeat structure through a single reaction, we constructed three germplasm lines of forward repeat, reverse repeat and intensive repeat for sequencing (FIG. 5). After interrupting the three germplasm lines with g-tube, we sequenced them with PacBio. After reverse repeat sequencing, the average sequencing depth was 250, and some bases had mutation, deletion, and insertion rates of less than 5%, which could be determined to be introduced by the PacBio sequencing machine (Figure 6). Such SNPS are equivalent to the low peaks of sanger sequencing and can be ignored. The average depth of forward repeat sequencing was 5000, and no deletion was found at the position of sequence repetition, indicating a good sequencing result. However, in the 621bp-624bp deletion occurred, the deletion rate was 60%. This sequence can therefore be determined to be a mixed clone (Figure 7). After intensive repeat sequencing, only some bases had mutations, deletions, and insertion rates below 5%, which could be judged to have been introduced by the PacBio sequencing machine (Figure 8).

**Figure 5.**
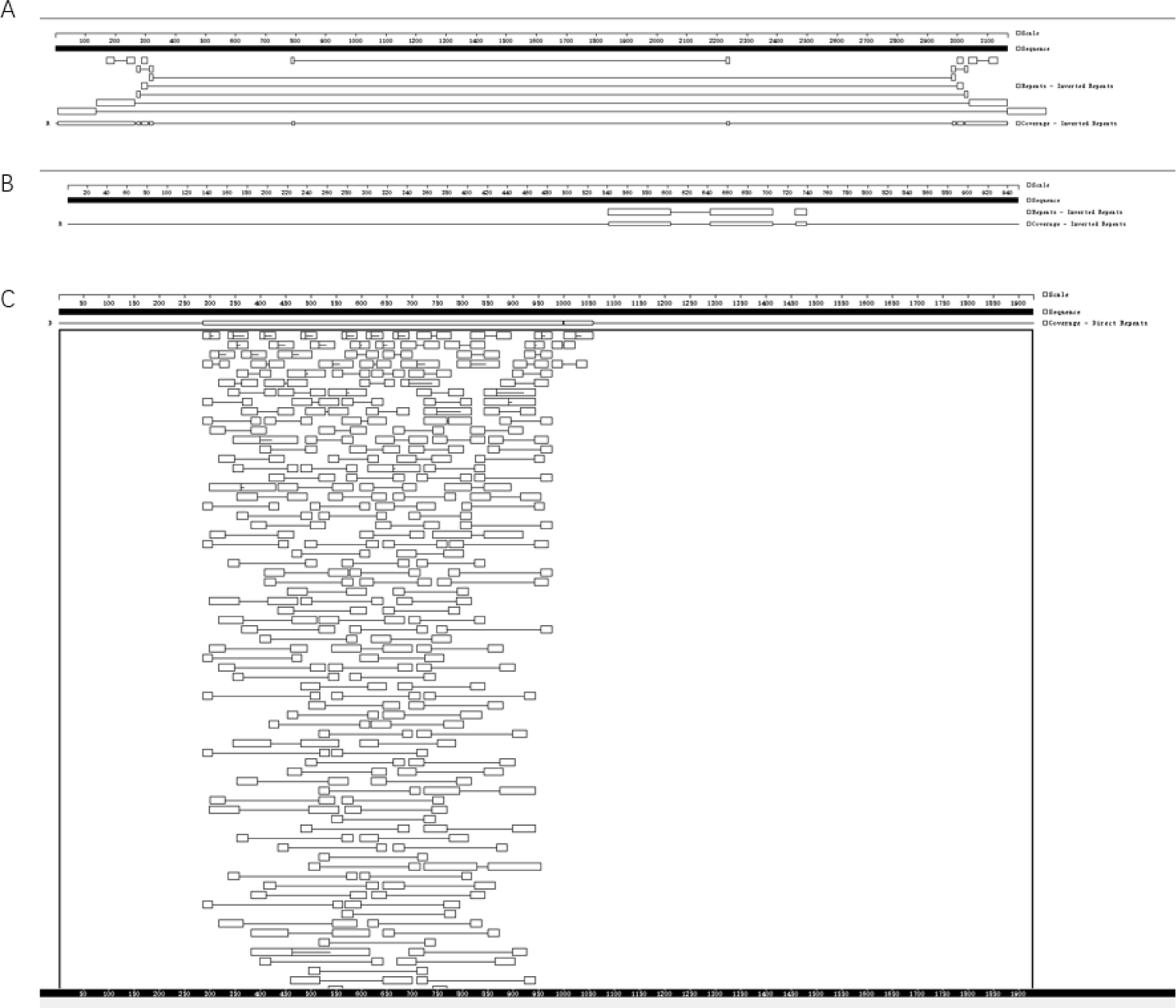
Sequence structure analysis of three repeating plasmids. Gene Quest software was used to analyze the series structure. (A) Reverse repeat sequence structure analysis. There are two reverse repeats of 129bp length at either end of the sequence. (B) Forward repetitive structural analysis. In the middle of the sequence are two forward repeats of length 29bp. (C) Intensive repeat sequence structural analysis. This sequence contains many repetitive structures larger than 20bp, mainly concentrated in 285bp-998bp, with a length of 713bp.

**Figure 6.**
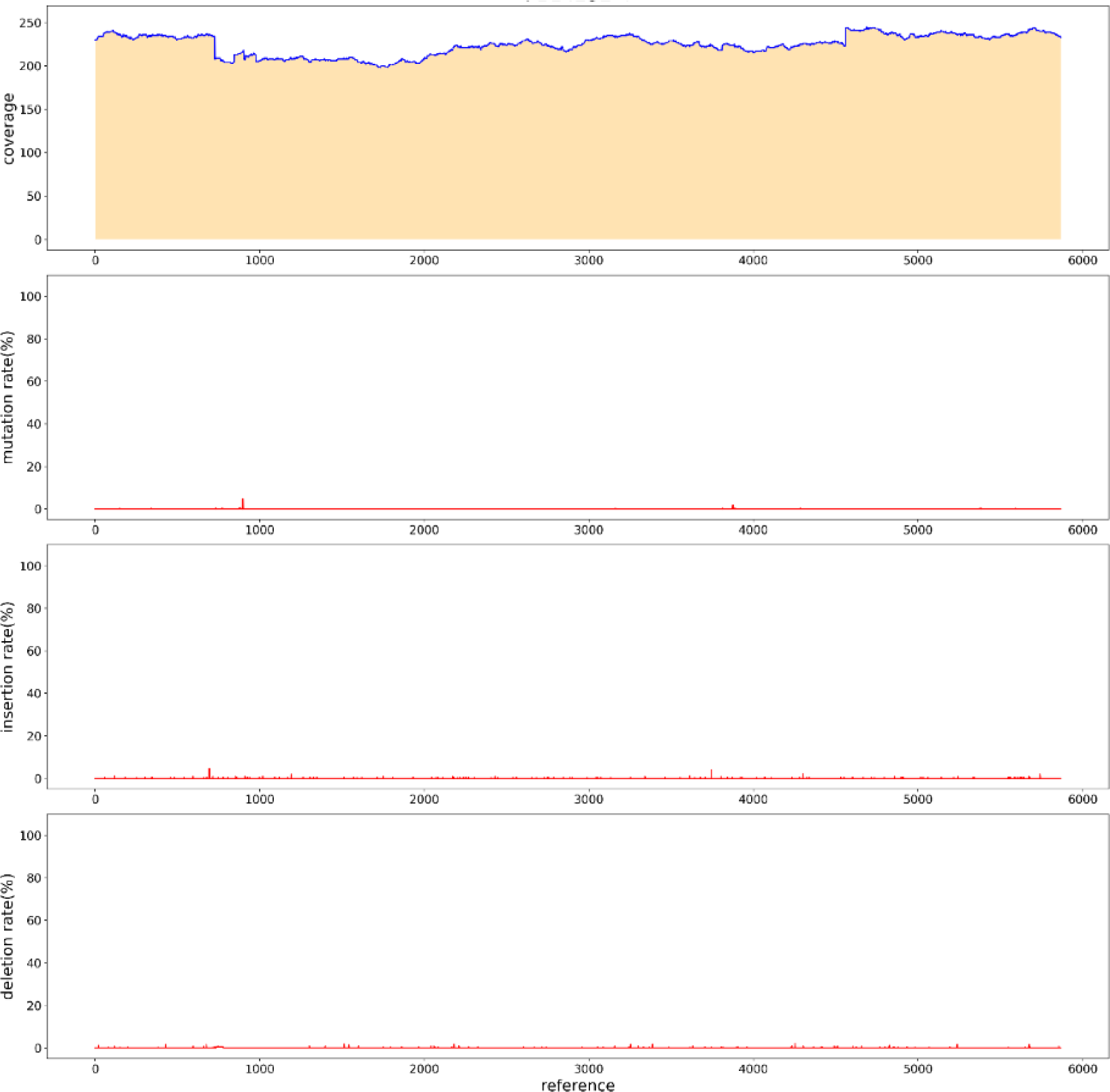
Reverse repeat sequence PacBio sequencing results. The sequencing depth was 230 with no mutations or insertions.

**Figure 7.**
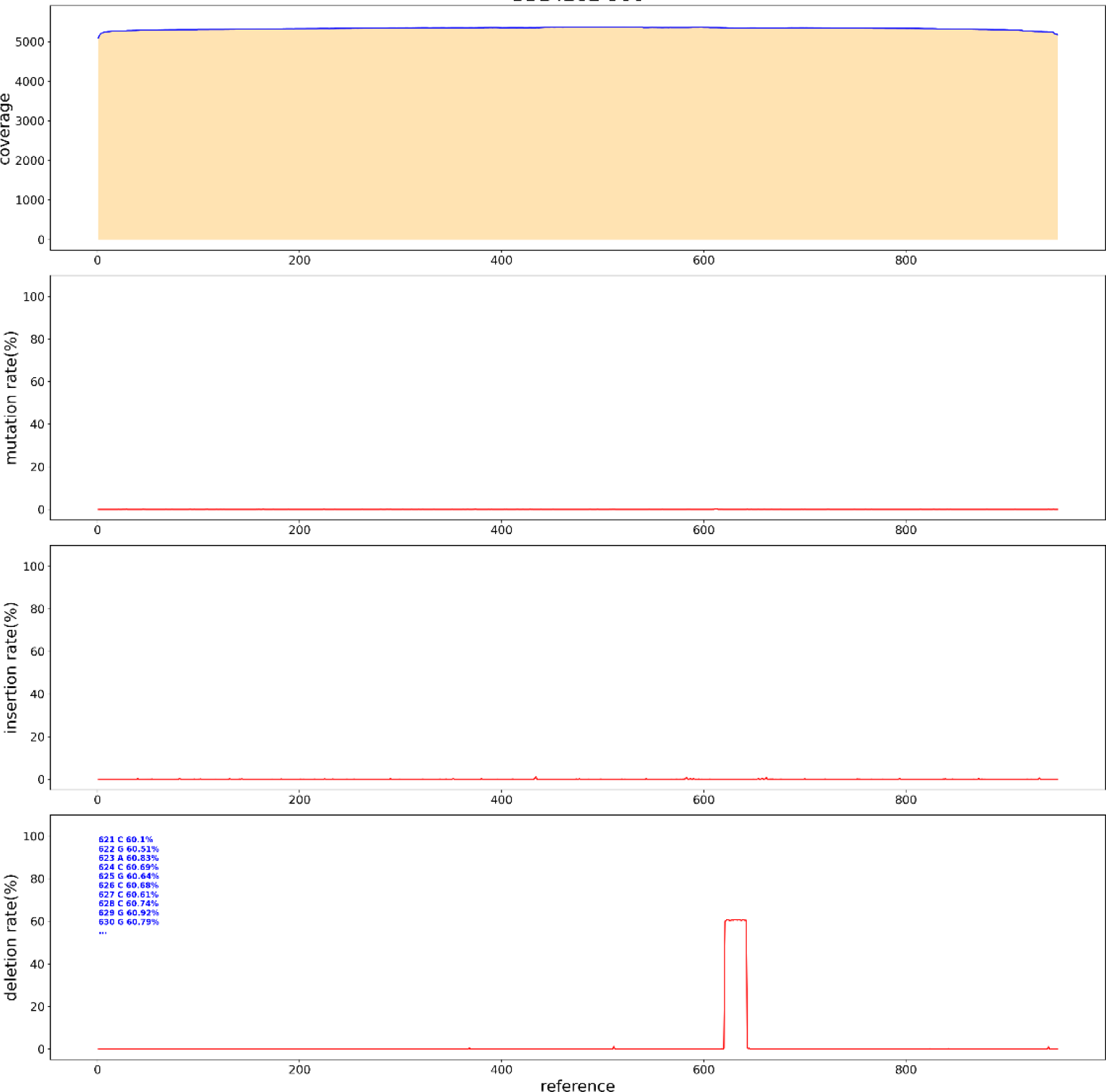
PacBio sequencing results of forward repeat sequence. The sequencing depth was 5000 with no mutations or insertions. Deletion occurred at 621bp-624bp, and the deletion rate was 60%.

**Figure 8.**
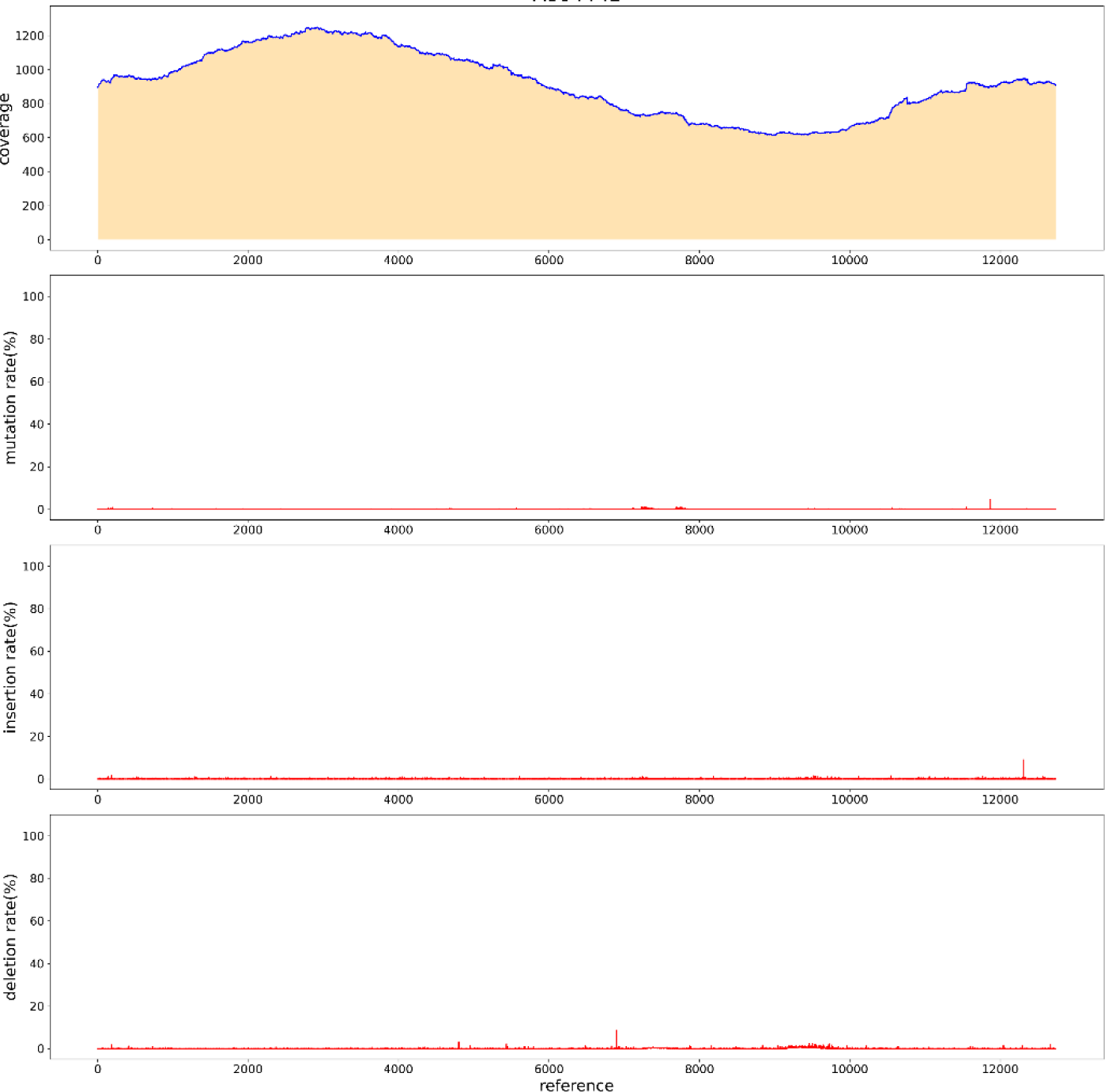
Results of PacBio sequencing of dense repeat sequences. The sequencing depth was 800 with no mutations, deletions, or insertions.

### High GC structure sequencing by PacBio

Sanger sequencing requires the design of sequencing primers, and the level of GC content affects the results of sanger sequencing. PacBio sequencing does not need to design sequencing primers to detect the whole sequence, which reduces the influence of GC on sequencing. To verify that PacBio could measure GC plasmid elevation, we loaded fragments with a GC value of 68% into pUC vectors, and the overall GC content was 53% (Figure 9). Interrupted with g-tube and sequenced with PacBio. BI analysis showed that the entire sequence was covered by reads, indicating that the entire sequence could be tested (Figure 10). Only some of the bases had mutations, deletions, and insertion rates of less than 5%, and could be judged to have been introduced by the PacBio sequencing machine (Figure 11).

**Figure 9.**
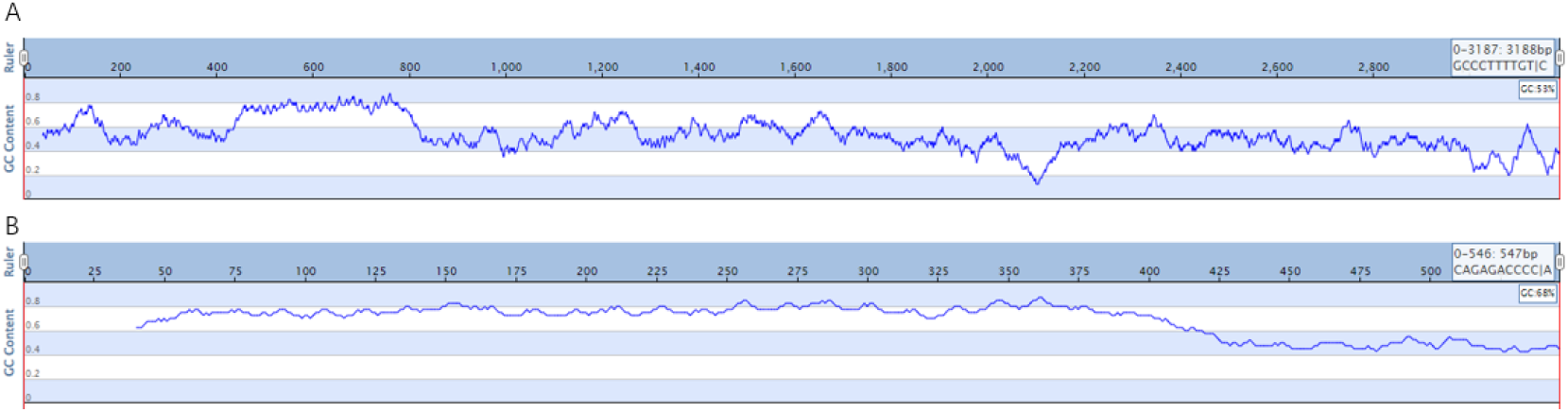
GC content analysis. (A) GC content analysis of whole sequence. (B) Insertion sequence GC content analysis.

**Figure 10.**
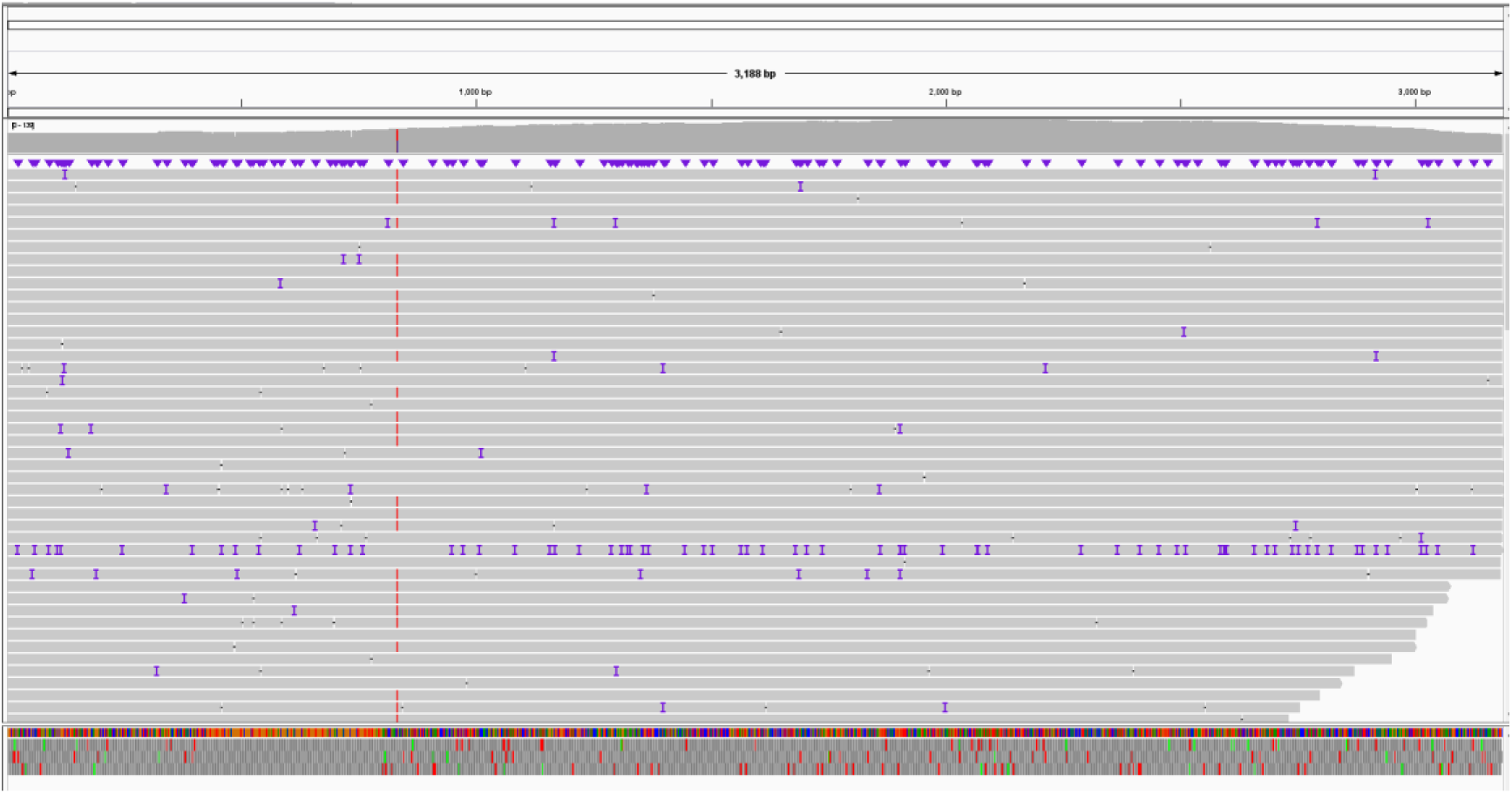
Distribution of sequence reads. IGV software was used to determine the distribution of reads in the whole sequence.

**Figure 11.**
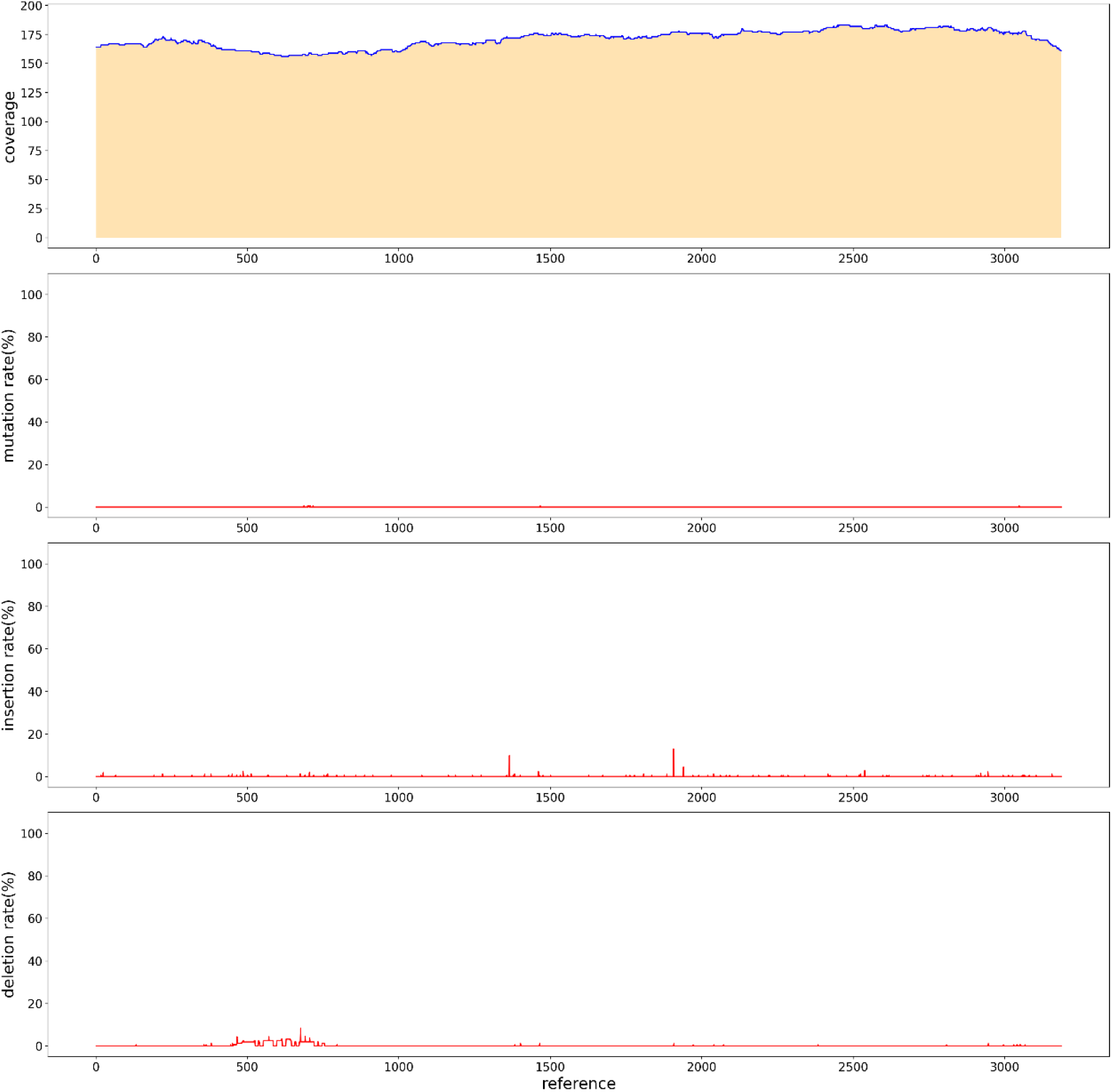
PacBio sequencing results of high GC sequence. The sequencing depth was 160 with no mutations, deletions, or insertions.

### Low-copy plasmids sequencing by PacBio

For some low-copy plasmids, lower concentration affected their sequencing results. Similarly, in PacBio sequencing, the amount of plasmid input affects the output of PacBio sequencing data. In order to increase the output of PacBio data, we measured whether the number of CCS increased by rolling loop replication before and after building the library. The first group of reactions: the library was built first, and then the rolling replication was carried out after the construction. As shown in Table 2, the concentration increased significantly after the rolling ring replication. When 20ng plasmids were put into PacBio sequencing, it was found that there was no amount of data for roll-over replication after the completion of library construction, and the CCS number of plasmids without roll-over replication was also very low due to the low concentration of plasmids in library construction (Table 3). In the second group of reactions, roll-over replication was performed first, and the library was constructed. The concentration was shown in Table 4. It could be found that the concentration was doubled, and the same number of plasmids were put in for PacBio sequencing. It can be found that the CCS value increases greatly after the loop replication is built, which is sufficient for analysis (Table 5)and the sequencing was successful(FIG. 14). Therefore, when the plasmid extraction concentration is low, it is less than above, we can increase the plasmid concentration by rolling replication, and then interrupt and build a database by red-gute, and the data volume will be significantly increased.

**Table 2.**
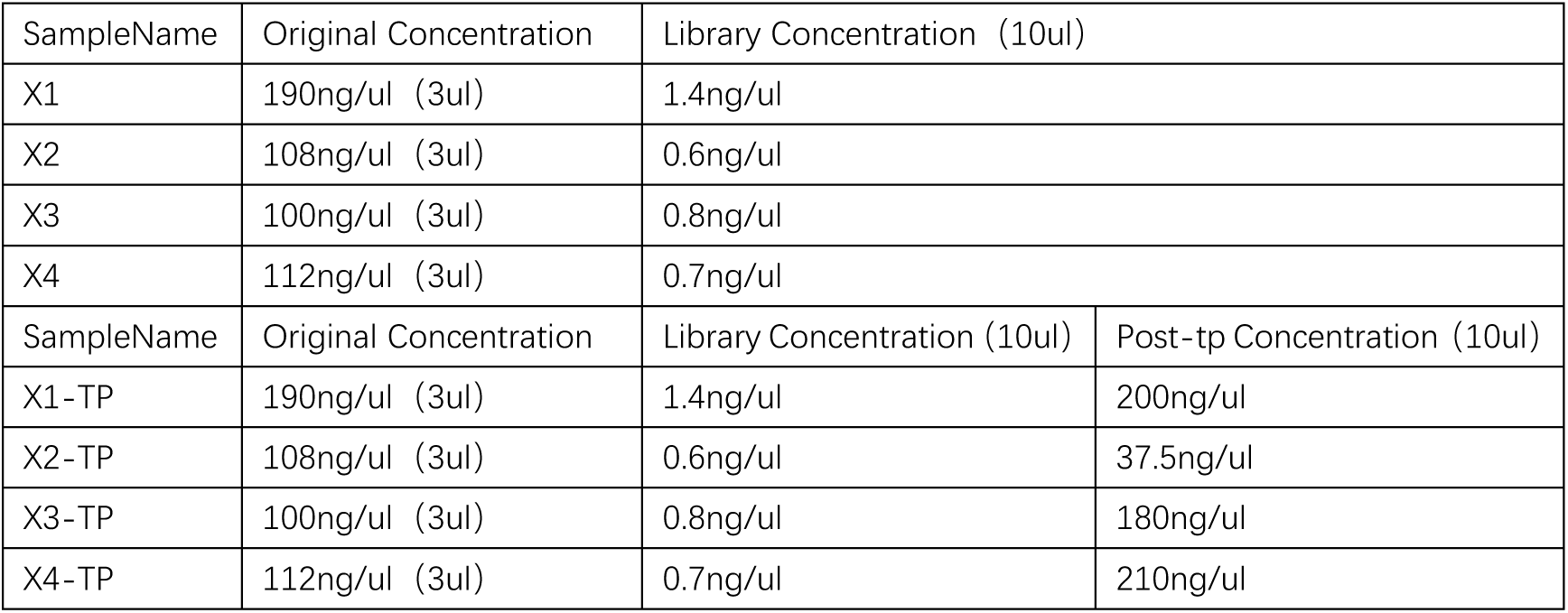
After the completion of library construction, roll out replication and plasmid concentration.

**Table 3.** Amount of Pacbio data for roll-over replication after database construction.

| SampleName | TotalReads | MappedReads | Percent(%) | UnmappedReads | Percent(%) |
| --- | --- | --- | --- | --- | --- |
| X1 | 6 | 2 | 33.33 | 4 | 66.67 |
| X1-TP | 1 | 1 | 100 | 0 | 0 |
| X2-59 | 9 | 8 | 88.89 | 1 | 11.11 |
| X2-TP | 0 | 0 | 100 | 0 | 0 |
| X3 | 9 | 4 | 44.44 | 5 | 55.56 |
| X3-TP | 0 | 0 | 100 | 0 | 0 |
| X4 | 9 | 4 | 44.44 | 5 | 55.56 |
| X4-TP | 0 | 0 | 100 | 0 | 0 |

**Table 4.** library construction after loop replication, plasmid concentration.

| SampleName | Original Concentration | Library Concentration (10ul) |  |
| --- | --- | --- | --- |
| X5 | 95ng/ul (6ul) | 3.5ng/ul |  |
| X6 | 209ng/ul (6ul) | 4.8ng/ul |  |
| X7 | 200ng/ul (6ul) | 3.7ng/ul |  |
| SampleName | Original Concentration | Post-tp Concentration | Library Concentration (10ul) |
| X5 | 95ng/ul (6ul) | 200ng/ul (70ul) | 6.8ng/ul |
| X6 | 209ng/ul (6ul) | 210ng/ul (70ul) | 10.2ng/ul |
| X7 | 200ng/ul (6ul) | 190ng/ul (70ul) | 7.6ng/ul |

**Table 5.** Amount of Pacbio data for database construction after rolling ring replication.

| SampleName | TotalReads | MappedReads | Percent(%) | UnmappedReads | Percent(%) | Coverage |
| --- | --- | --- | --- | --- | --- | --- |
| X5 | 57 | 44 | 77.19 | 13 | 22.81 | 11 |
| X5-TP | 926 | 848 | 91.58 | 78 | 8.42 | 282 |
| X6 | 38 | 24 | 63.16 | 14 | 36.84 | 4 |
| X6-TP | 1012 | 955 | 94.37 | 57 | 5.63 | 375 |
| X7 | 106 | 86 | 81.13 | 20 | 18.87 | 27 |
| X7-TP | 894 | 838 | 93.74 | 56 | 6.26 | 305 |

**Figure 12.**
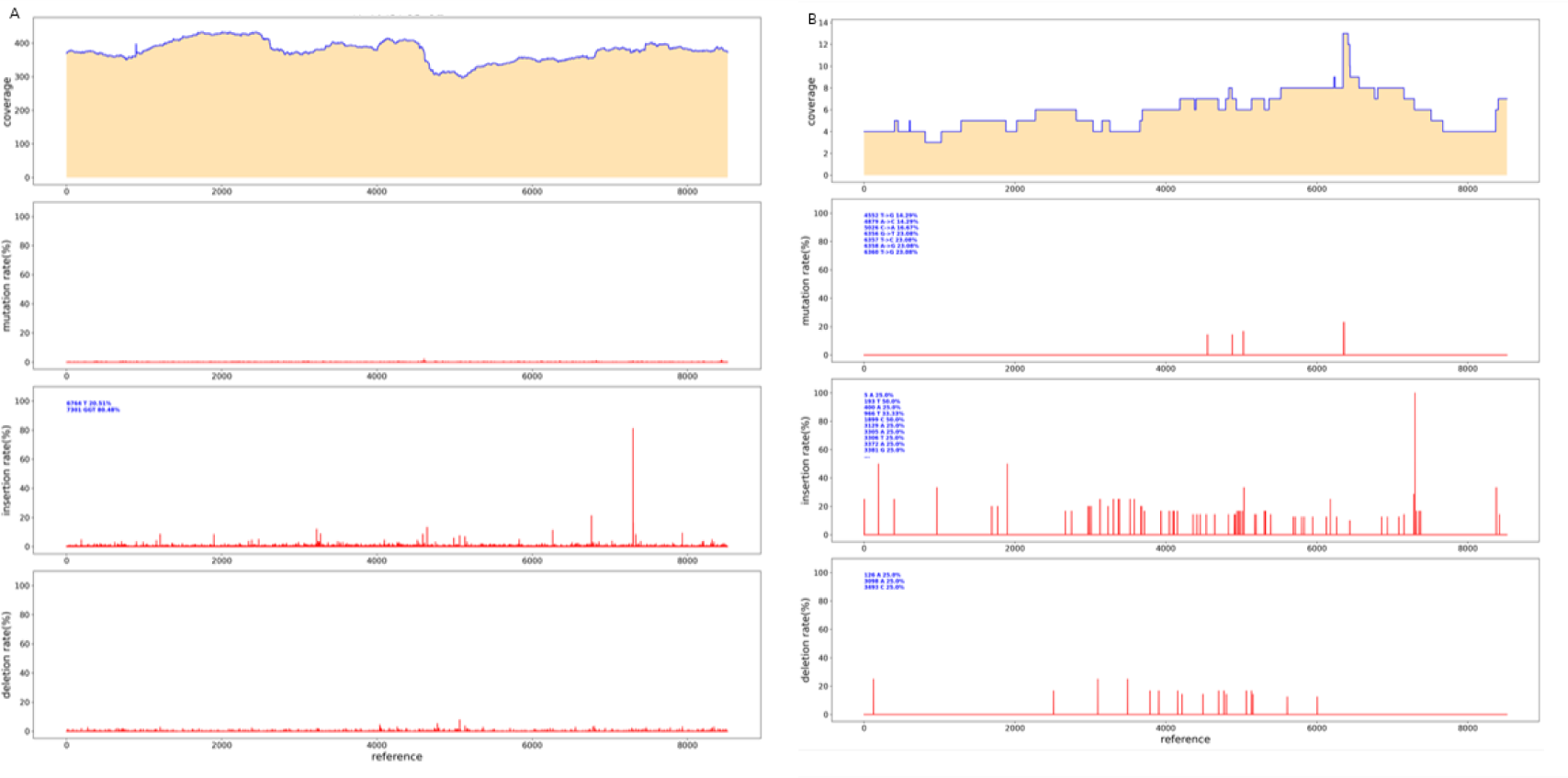
Pacbio sequencing results.(A) After roll out replication, Pacbio sequencing results.(B) Untreated plasmid, Pacbio sequencing results

### De novo assembly for Parameter less sequences

For parameter less sequences, the main method of Sanger sequencing is to randomly design universal primers. If the sequence is detected, the primers will continue to be designed according to the detected sequence; if no sequence is detected, the universal primers will continue to be designed until a sequence is detected by the universal sequencing primer. Using sanger to measure parameter less sequences is costly, time-consuming, and stochastic. To verify whether PacBio can measure parameter less sequences, we randomly took a plasmid and used rubies to break the library. Through BI analysis, we assembled A plasmid sequence of 11436bp (FIG. 13), the sequencing depth was 3700ccs, only at the position of 10546bp, an A base was inserted, and the insertion rate was 16.83% (FIG. 14). sanger sequencing can be used in the later stage to confirm its correctness. Thus, it can be shown that PacBio can measure parameter less sequences, and BI can be spliced into a whole sequence through analysis, which is short in time and low in cost.

**Figure 13.**
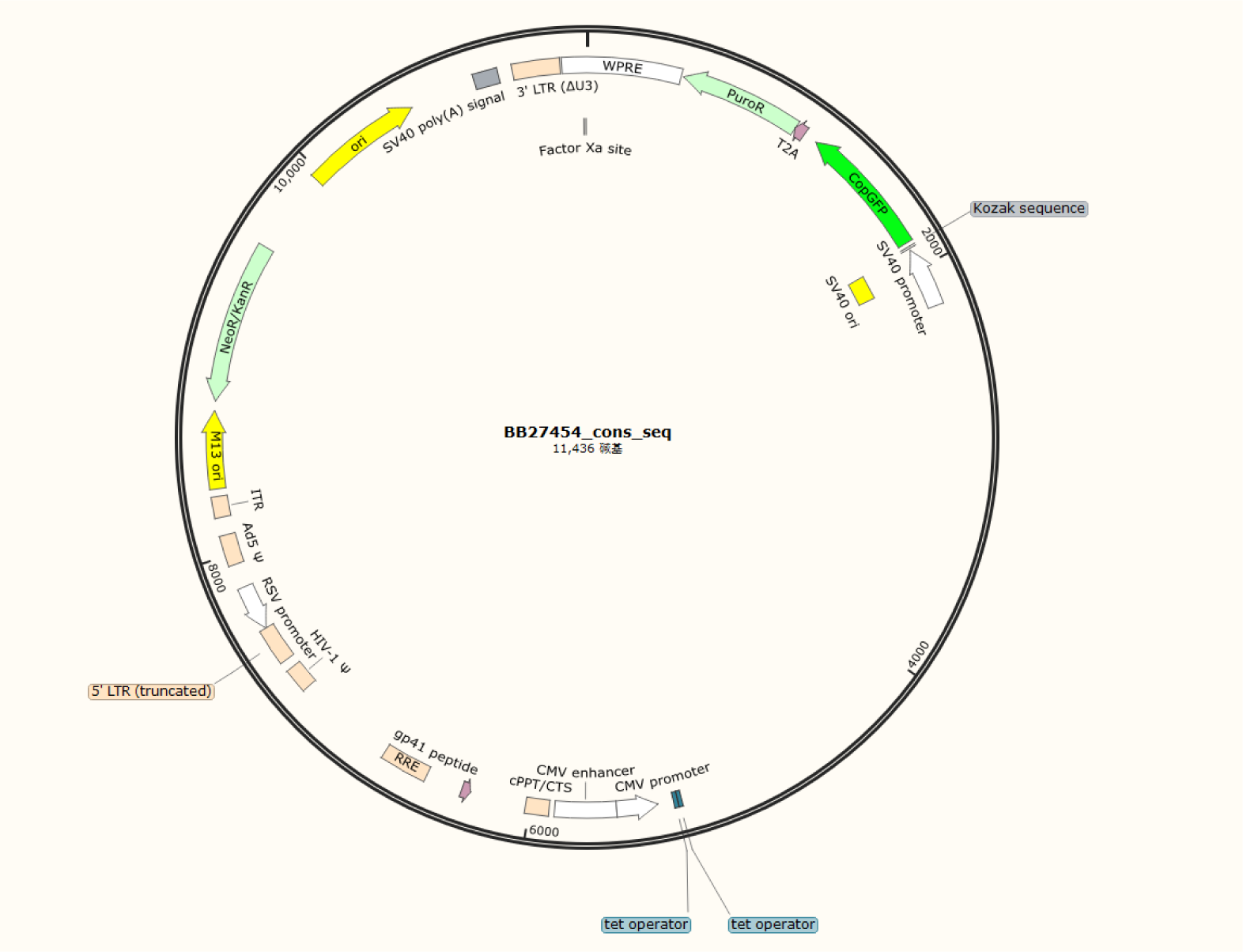
Map of PacBio sequencing results without reference sequence. The total length is 11436bp, and the whole sequence is clear.

**Figure 14.**
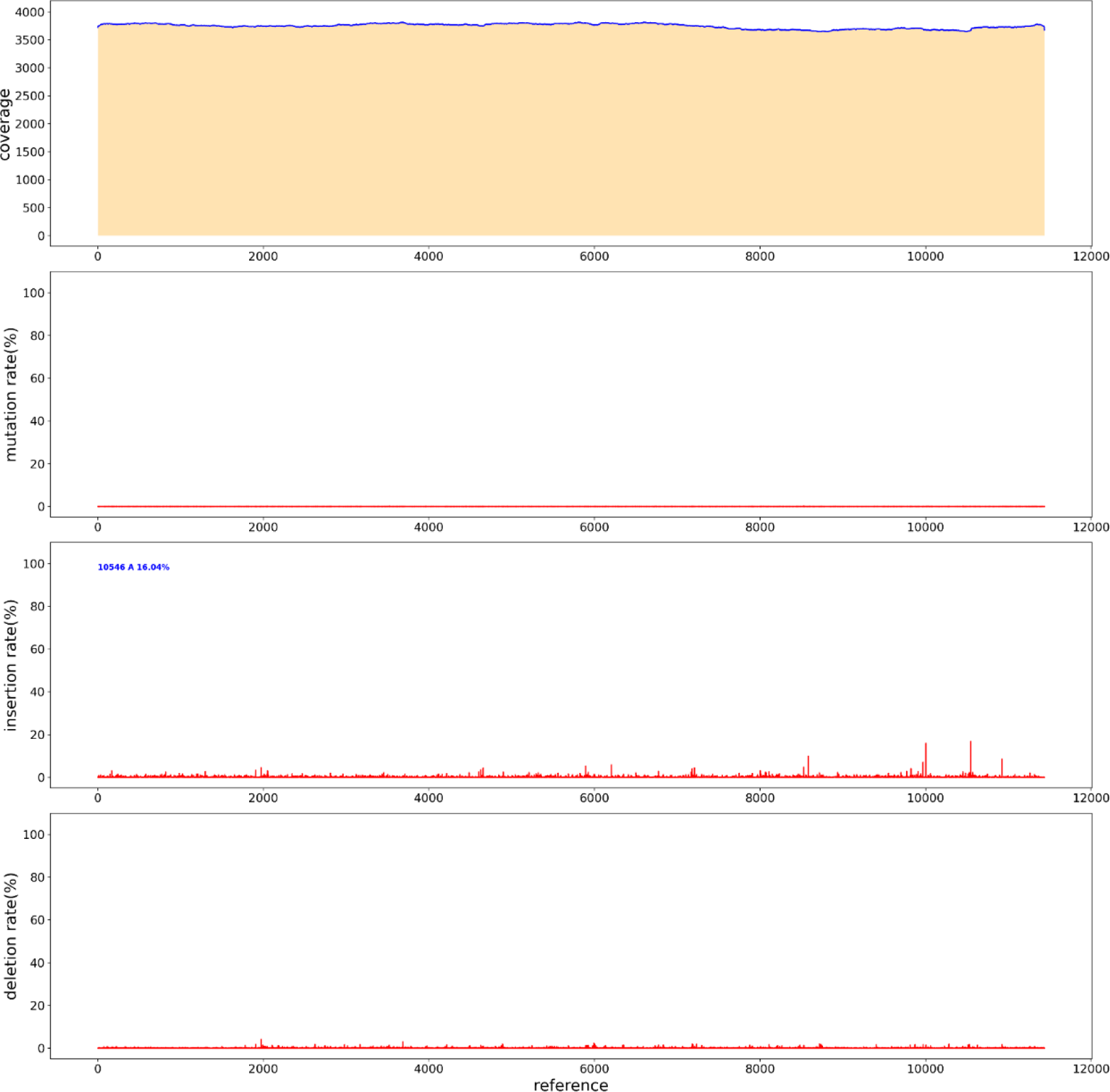
Results of PacBio sequencing without reference sequence. The sequencing depth was 3700, only at the position of 10546bp, an A base was inserted, and the insertion rate was 16.83%

### Simplify the database building process

In order to simplify the process of library construction, A restriction enzyme at the flat end was selected, and the carrier was enzymically cut first, and the product after the enzyme cut was directly connected to the adapter. The two steps of adding A-tail and repairing DNA fragments were omitted. Redesigned the adapter sequence, in addition to the original “T” bases, sequence is: / 5 phos/ATCTGATAGAGTGTGTATCTCTCTCTTTTCCTCCTCCTCCGTTGTTGTTGTTGAGAGAGATACACACTCTATCAGAT. After the plasmid digested by NruI enzyme and recovered, the plasmid was directly added by adapter with T4 ligase. The sequencing results of PacBio showed (FIG. 15) that the average sequencing depth was 1200, and no defects were found in the sequence, indicating a good sequencing result.

**Figure 15.**
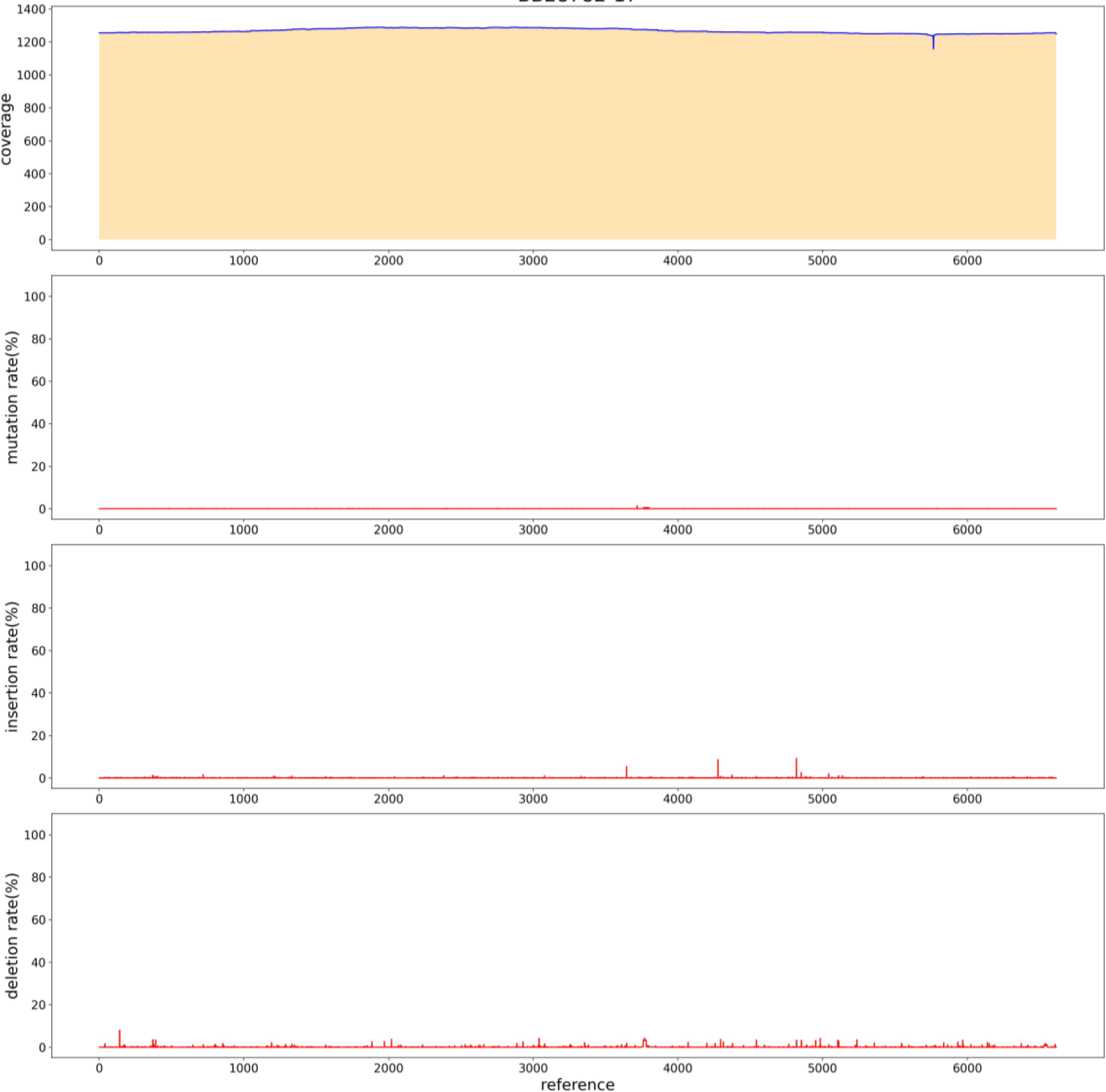
The result of PacBio sequencing after simplifying the database construction process. The sequencing depth was 1200 with no mutations or insertions.

## Methods

### Plasmid construction

Primers were designed to synthesize the target fragment, and the target sequence (GENEWIZ biotech GsmartI Pfu DNA PolymeraseDP1701S) was amplified by PCR. Then the vector was linearized by restriction endonuclease. Then T4 ligase (GENEWIZ biotechT4 DNA LigaseCR1201S) was used to connect the target fragment to the carrier. Competent Cells was obtained by DH5α Chemically Competent Cell (Weidibio DH5α Chemically Competent Cell 1001S). The single clone was selected and verified by PCR (Gsmart Taq DNA PolymeraseDP1702S of Suzhou GENEWIZ biotech Co., LTD.), and the correct positive clone was obtained. Plasmid extraction (AxyPrep plasmid DNA small dose kit AP-MN-P-50) for PacBio sequencing.

### PacBio library preparation

The plasmid was interrupted with g-TUBE (Covaris g-TUBE520079), and the plasmid was repaired with New England Biolabs PreCR^®^ Repair MixM0309L. 1μL of 5: 1dATP (Vazyme dATPP034-01)/dNTP (Vazyme dNTP Mix P031-01); 0.5μL T4 PNK (Vazyme T4 Polynucleotide KinaseN102-01); 2μL T4 DNA Polymerase (Vazyme T4 DNA polymerase N101-01); 0.5μL rTaq (Takara TaKaRa Taq™R001A) plus A tail for connection, 8.0 μLPrD (aladdin PrD P103430) at the end; 10.0 μL50% PEG-8000 (solarbio PEG 8000 25322-68-3); 4.0μLT4 DNA Ligase (Vazyme T4 DNA LigaseC301-01) was connected to the interrupted plasmid, the library was constructed, and the fragment length was tested using Agilent 2100 for quality inspection. The constructed SMRTbell library, combined with the upper primers and polymerase of the SMRTbell Express Template Prep Kit 2.0 kit, was added to the sequencing chip of the PacBio platform in a freely diffused manner for sequencing

### PacBio data analysis

The original disembarking data is first corrected by using ccs (4.2.0) [8] software for subread generated by the same zero-mode waveguide hole (ZMW) to obtain a high-quality CCS sequence. minimap2 (version 2.15-r905)[9] software is used to compare the high-quality CCS sequence to the reference sequence. Finally, samtools(Version: 1.9) [10] software was used for quality control of the comparison results. Specific quality control parameters include the minimum cycle sequencing times, the minimum prediction accuracy, the minimum insertion fragment length and the maximum insertion fragment length, and the set values of the above parameters are 5,90,1 400 and 1 800, respectively. After quality control, the qualified sequences were divided into different samples according to barcode sequence. Meanwhile, barcode and primer information of each sequence were deleted for subsequent analysis.

## Discussion

The maximum sequence length of Sanger sequencing was about 1 kb. It had high requirements on the total amount of DNA, and generally cloned the target DNA sequence and connected the vector[21], and then amplified by prokaryotic E. coli (at that time, the genome De novo was sequenced by BAC library), which was long and time-consuming. Illumina is a major supplier in the sequencing market due to its low cost, high speed, and high yield. The Illumina sequencing platform is widely applicable[22], so NGS has been widely used to explore various areas of genomics, including oncology, microbiology, environmental genomics, metagenomics and medical, environmental, and agricultural research, and its wide range of applications[23]. The disadvantage is also gradually prominent, that is, the reading length of second-generation sequencing (represented by Illumina) is still an important bottleneck in biological research, which limits the accuracy of many biological studies, especially in genome assembly research. The use of short-read long sequencing in segmental duplication, structural variations (SV), or paralogue analysis may result in a large number of false positives. Similar to Sanger sequencing and NGS sequencing [24], PacBio sequencing also uses synthesis side by side sequencing, using one DNA strand as a template and synthesizing another strand through DNA polymerase.

PacBio has upgraded CCS Sequencing mode for Long read High Fidelity (HiFi) 15 kb reads, Circular Consensus Sequencing (CCS) read: A circular consistency sequence, which is generated by comparing subreads from a single ZMW [25]. The generated CCS reads require at least three rounds of reading of subreads from inserted fragments using CCS algorithm, and the accuracy of a single CCS read can reach more than 99%. We tested both 50kb and 32kb plasmids, and the correct sequence was detected [26]. The CCS model we used needs to be tested on the dosage to achieve higher coverage. During the construction of the library by Pacbio, we connected the joint to the carrier in the way of connection. The whole process did not involve PCR process, and it could detect regions rich in AT or GC, highly repeated sequences, palindromic sequences, etc., which would not cause large deviation of GC, and would not cause uneven coverage of all fragments due to the specificity of PCR [27]. Through pacbio sequencing, we solved the difficult Sanger of recombinant adeno-associated viruses with high GC, multi-structure, long sequence, genome and repeat sequence. Since the principle of Sanger sequencing is different from that of pacbio sequencing, Sanger needs to design corresponding primers according to the reference sequence and then sequence. While in the process of building the library of pacbio sequencing, fragments need to be broken and loaded into the adapter, which contains primers and index used for sequencing. Therefore, it is not necessary to redesign sequencing primers, so that parameterless sequences can be measured [28]. Sanger sequencing requires the use of a large number of plasmids, so it is necessary to greatly reserve plasmids for sequencing in the synthesis process. We conducted rolling loop replication of the plasmids before sequencing by pacbio, reducing the use of plasmids and the cost of synthesis. At the same time, the sequencing joint was connected by the method of enzyme digestion, which reduced the sequencing time.

Notably, the ability of PacBio sequencing to process single-stranded DNA input should not be overstated, as it still favors double-stranded templates, and we observed read length compression in PacBio, where the mapped carrier genome read length was 2-3% shorter than expected [29]. This phenomenon was found to be related to the high frequency gaps/omissions found in the read length, rather than due to poor processing power. Since the genome length defined by pacbio read length is not very reliable, we recommend defining truncation by aligning the start and end of the read length [30]. Although this strategy does not necessarily yield information about the structure of the carrier genome, analyzing the PacBio reading results by the starting and ending positions of the alignment can still provide information about the functional regions across. This information can be used to infer the percentage of the functional genome during preparation.

## ACKNOWLEDGMENTS

We thank members of Azenta for their technical support in genome sequencing.

## Supplement

SI Tabl:Large DNA fragment pacbio sequencing results

S2 Tabl:PolyA structure pacbio sequencing results

S3 Tabl:PolyC structure pacbio sequencing results

S4 Tabl:Poly structure pacbio sequencing results

S5 Tabl:Forward repeated structure pacbio sequencing results

S6 Tabl:Intensive repeated structure pacbio sequencing results

S7 Tabl:high GC plasmid pacbio sequencing results

S8 Tabl:No parameter sequence, pacbio sequencing result

S9 Tabl:The result of pacbio sequencing after simplifying the database construction process.

S10-17 Tabl:Pacbio sequencing results of X1-X4.

S18-23 Tabl:Pacbio sequencing results of X5-X7.

## References

1. Rhoads A, Au KF. PacBio sequencing and its applications. Genom Proteome Bioinf. 2015;13(5):278–289. doi: 10.1016/j.gpb.2015.08.002.

2. Bayega A, Fahiminiya S, Oikonomopoulos S, et al. Current and future methods for mRNA analysis: a drive toward single molecule sequencing//gene expression analysis. New York: Humana Press; 2018. pp. 209–241.

3. Schadt E.E., Turner S., Kasarskis A. A window into third-generation sequencing. Hum Mol Genet. 2010;19:R227–R240.

4. Pacific Biosciences. Media Kit, <http://www.pacb.com/company/news-events/media-resources/page/3/> (May 19, 2015, date last accessed).

5. Eid J., Fehr A., Gray J., et al. Real-time DNA sequencing from single polymerase molecules. Science. 2009;323:133–138.

6. Levene MJ, Korlach J, Turner SW, et al. 2003. Zero-mode waveguides for single-molecule analysis at high concentrations. Science, 299(5607): 682–686.

7. Roberts RJ, Carneiro MO, Schatz MC 2013. The advantages of SMRT sequencing. Genome Biol, 14(7): 405.

8. Gorrieri R, Versari C. CCS: A Calculus of Communicating Systems[J]. Springer International Publishing, 2015.

9. Heng. Minimap2: pairwise alignment for nucleotide sequences[J]. Bioinformatics, 2018.

10. Danecek P, Bonfield J K, Liddle J, et al. Twelve years of SAMtools and BCFtools[J]. GigaScience, 2021, 10(2).

11. Wang B, Tseng E, Regulski M, et al,. Unveiling the complexity of the maize transcriptome by single-molecule long-read sequencing. Nat Commun. 2016;7:11708. doi: 10.1038/ncomms11708.

12. Schmidt MHW, Vogel A, Denton AK, et al,. De novo assembly of a new Solanum pennellii accession using nanopore sequencing. Plant Cell. 2017;29(10):2336–2348. doi: 10.1105/tpc.17.00521.

13. Chao Y, Yuan J, Li S,, et al. Analysis of transcripts and splice isoforms in red clover (Trifolium pratense L.) by single-molecule long-read sequencing. BMC Plant Biol. 2018;18(1):300. doi: 10.1186/s12870-018-1534-8.

14. Finn RD, Bateman A, Clements J, et al,. Pfam: the protein families database. Nucleic Acids Res. 2014;42:D222–D230. doi: 10.1093/nar/gkt1223.

15. Wenger AM, Peluso P, Rowell WJ, et al. Accurate circular consensus long-read sequencing improves variant detection and assembly of a human genome. Nat Biotechnol. 2019;3710:1155–1162. doi: 10.1038/s41587-019-0217-9.

16. PacBio. Technical overview: HiFi library preparation using SMRTbell express template prep kit 2.0 for de novo assembly and variant detection applications. 2021. HiFi Library Preparation Using SMRTbell Express TPK 2.0 for De Novo Assembly and Variant Detection.

17. Hon T, Mars K, Young G, et al. Highly accurate long-read HiFi sequencing data for five complex genomes. Sci Data. 2020;7:399. doi: 10.1038/s41597-020-00743-4.

18. Nishii K, Möller M, Hart M. High molecular weight DNA extraction for long-read sequencing v.1. Protocol.io. 2022 doi: 10.17504/protocols.io.bempjc5n.

19. van Rengs WMJ, Schmidt MH, Effgen S, Le DB, Wang Y, et al. A chromosome scale tomato genome built from complementary PacBio and Nanopore sequences alone reveals extensive linkage drag during breeding. Plant J. 2022;110:572–588. doi: 10.1111/tpj.15690.

20. De Coster W, D’Hert S, Schultz DT, Cruts M, Van Broeckhoven C. NanoPack: visualizing and processing long-read sequencing data. Bioinformatics. 2018;34:2666–2669. doi: 10.1093/bioinformatics/bty149.

21. Chin CS, Alexander DH, Marks P, et al. Nonhybrid, finished microbial genome assemblies from long-read SMRT sequencing data. Nat Methods. 2013;10(6):563–569. doi: 10.1038/nmeth.2474.

22. Otto TD, Sanders M, Berriman M, et al. Iterative Correction of Reference Nucleotides (iCORN) using second generation sequencing technology. Bioinformatics. 2010;26(14):1704–1707. doi: 10.1093/bioinformatics/btq269.

23. Simpson JT, Durbin R. Efficient de novo assembly of large genomes using compressed data structures. Genome Res. 2012;22(3):549–556. doi: 10.1101/gr.126953.111.

24. Rhoads A., Au K. F. (2015). PacBio sequencing and its applications. Genomics Proteomics Bioinformatics 13 278–289. 10.1016/j.gpb.2015.08.002

25. Xia Y., Yang C., Zhang T. (2018). Microbial effects of part-stream low-frequency ultrasonic pretreatment on sludge anaerobic digestion as revealed by high-throughput sequencing-based metagenomics and metatranscriptomics. Biotechnol. Biofuels 11:47.

26. Ye C., Hill C. M., Wu S., Ruan J., et al. DBG2OLC: efficient assembly of large genomes using long erroneous reads of the third generation sequencing technologies. Sci. Rep. 6:31900.

27. França LTC, Carrilho E, Kist TBL. A review of DNA sequencing techniques. Q Rev Biophys. 2002;35:169–200. doi: 10.1017/S0033583502003797.

28. Maricic T, Whitten M, Pääbo S. Multiplexed DNA sequence capture of mitochondrial genomes using PCR products. PLoS ONE. 2010;5:e14004. doi: 10.1371/journal.pone.0014004.

29. Cruaud P, Rasplus JY, Rodriguez LJ, Cruaud A. High-throughput sequencing of multiple amplicons for barcoding and integrative taxonomy. Sci Rep 2017 71. 2017;7:1–12. doi: 10.1038/srep41948.

30. Shokralla S, Gibson JF, Nikbakht H, et al. Next-generation DNA barcoding: using next-generation sequencing to enhance and accelerate DNA barcode capture from single specimens. Mol Ecol Resour. 2014;14:892–901. doi: 10.1111/1755-0998.12236.

